# Reconstructing complex lineage trees from scRNA-seq data using MERLoT

**DOI:** 10.1101/261768

**Authors:** R. Gonzalo Parra, Nikolaos Papadopoulos, Laura Ahumada-Arranz, Jakob El Kholtei, Noah Mottelson, Yehor Horokhovsky, Barbara Treutlein, Johannes Soeding

## Abstract

Advances in single-cell transcriptomics techniques are revolutionizing studies of cellular differentiation and heterogeneity. Consequently, it becomes possible to track the trajectory of thousands of genes across the cellular lineage trees that represent the temporal emergence of cell types during dynamic processes. However, reconstruction of cellular lineage trees with more than a few cell fates has proved challenging. We present MERLoT (https://github.com/soedinglab/merlot), a flexible and user-friendly tool to reconstruct complex lineage trees from single-cell transcriptomics data and further impute temporal gene expression profiles along the reconstructed tree structures. We demonstrate MERLoT’s capabilities on various real cases and hundreds of simulated datasets.

## Background

Recent advances in single-cell sequencing techniques [1, 2, 3] permit to measure the expression profiles of tens of thousands of cells making ambitious projects like the single-cell transcriptional profiling of a whole organism [4] or the human cell atlas [5] possible. These efforts will better characterize the different cell types in multicellular organisms and their lineage relationships [6]. These advances also put within reach the question of how single cells develop into tissues, organs or entire organisms, one of the most fascinating and ambitious goals in biology that would also have wide-ranging consequences for the study of many human diseases.

It is critical to develop methods that can reliably reconstruct cellular lineage trees that reflect the process by which mature cell types differentiate from progenitor cells. This is challenging due to the inherently high statistical noise levels in single cell transcriptomes, the high-dimensionality of gene expression space, and the strong non-linearities [6].

Different methods have been developed in the last years for inferring single-cell trajectories [7, 8]. Most of these methods first apply a manifold embedding in order to reduce the dimensionality of the problem and then implement various strategies for reconstructing the trajectory structure on it. Some tools are intended for linear topologies, while others aim to resolve bifurcations, multifurcations, or even complex trees with many internal branchpoints. The latter case has proven very challenging, and there is much room for improvement. Here we present MERLoT, a tool that can reconstruct highly complex tree topologies containing multiple cell types and bifurcations. MERLoT uses a lowdimensional embedding to locate the cellular lineage tree and then refines this structure in successive steps.

Different manifolds have been shown to be useful for the reconstruction of different lineage trees. MERLoT implements diffusion maps [9] as produced by the Destiny package [10] as the default method for dimensionality reduction. However, users can provide MERLoT with any low-dimensional space coordinates to perform the tree reconstruction. MERLoT explicitly models the tree structure, defining its endpoints, branchpoints, and locating a set of support nodes between these that act as local neighborhoods for cells. This model-based strategy gives insights into the temporal order of branching and the emergence of intermediary cell types. Once the lineage tree has been reconstructed in the low dimensional space, MERLoT is able to embed it back to the high dimensional gene expression space. The support nodes play a two-fold role in this step: they integrate the gene expression information of the cells assigned to them, and they inform the gene expression profiles of nearby support nodes. This reduces the overall noise levels, interpolates gene expression values for lowly sampled regions of the lineage tree and imputes missing expression values.

We show MERLoT’s performance on several real datasets, using different manifold embeddings, and on hundreds of simulated datasets. We generated a total of 1000 synthetic datasets with PROSSTT [11], divided into subsets of 100 simulations containing from 1 to 10 bifurcations. To the best of our knowledge, this benchmark dataset is the largest and more complete one up to date (available at http://wwwuser.gwdg.de/~compbiol/merlot/). We show that MERLoT outperforms other methods by producing a better classification of cells to the different branches that constitute the lineage trees. This is crucial when studying the progression of gene expression along the different trajectories in the tree, since a sub-optimal classification of cells mixing different cell types together leads to inaccurate imputation of gene expression time courses and impairs downstream analysis [3].

We repeated the benchmark with simulations generated by another tool, Splatter [12]. For more information, details about method performance, and divergence analysis of the simulations, please refer to the supplementary material.

MERLoT is implemented as an R package and publicly available at https://github.com/soedinglab/merlot. MER-LoT allows users to easily retrieve subpopulations of cells that belong to specific branches or belong to specific paths along the tree. It can also calculate pseudotime assignments, impute pseudotemporal gene expression profiles, or find genes that are differentially expressed on different tree segments.

## Results and discussion

### MERLoT’s workflow for lineage trees reconstruction and gene expression imputation

Given the matrix of expression values for all cells (Fig. 1A), MERLoT reconstructs lineage trees according to the following steps (for details see Methods): First, MERLoT applies a dimensionality reduction method to map the highdimensional expression vectors of cells to a low-dimensional space. Users can replace the default, diffusion maps, with the method of their choice. Second, MERLoT calculates a scaffold tree in the low-dimensional space combining the Dijkstra’s shortest path [13] and Neighbor Joining [14] algorithms to define the location of endpoints, branchpoints and their connectivity (Fig. 1B). The scaffold tree is used as initialization for an Elastic Principal Tree (EPT) [15]. The EPT smoothes the scaffold via an optimization procedure that places a user-defined number of support nodes between endpoints and their corresponding branchpoints interpolating the density of cells in the low-dimensional space (Fig. 1C). Once the low-dimensional tree is optimized, an initial pseudotime to is assigned to the user-specified tree root. The pseudotime of each cell is then proportional to its distance from the root along the tree structure (Fig. 1D).

**Figure 1:**
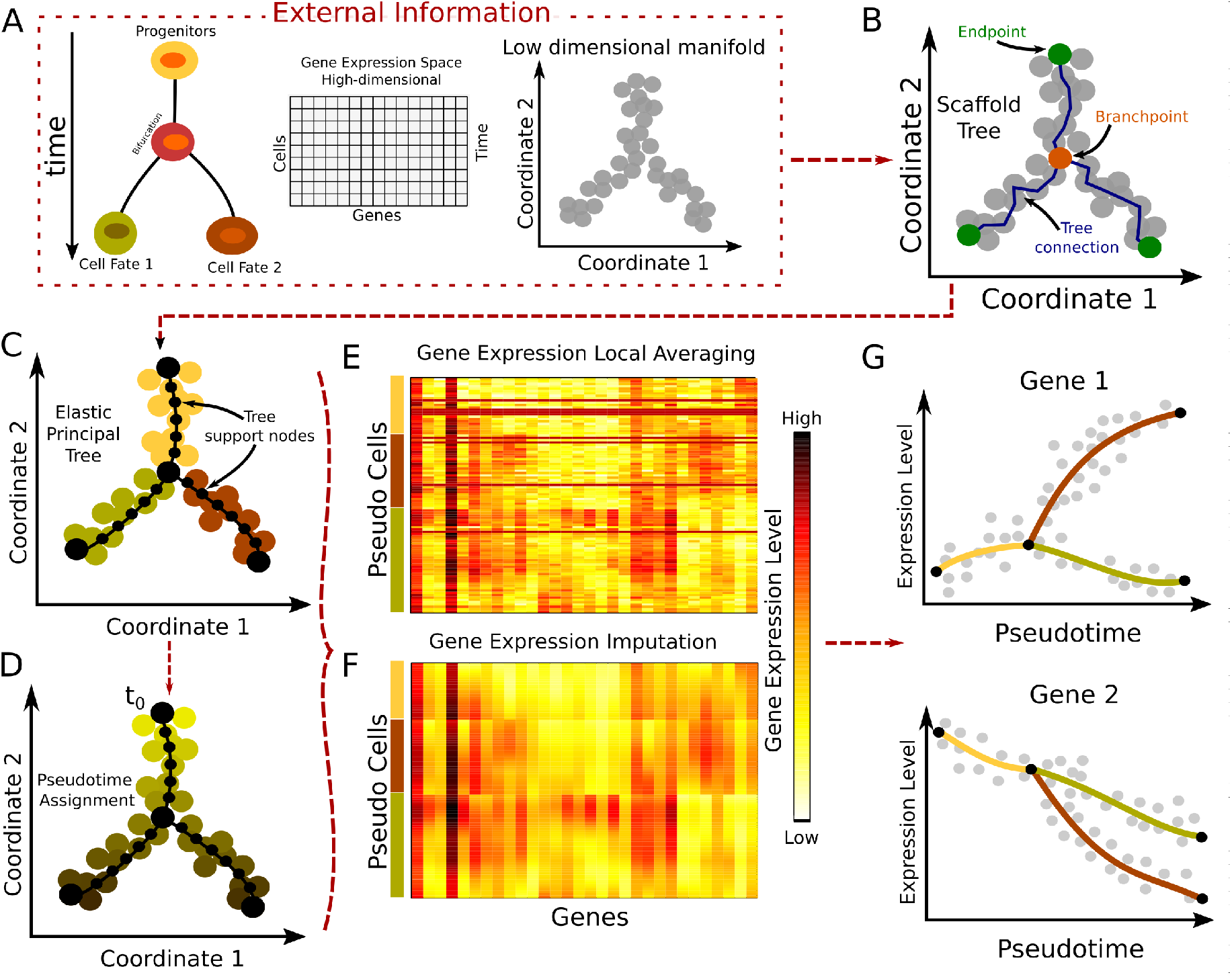
MERLoT’s workflow. **(A)** Input to MERLoT is a gene expression matrix sampled from a dynamic process in which several cell types are present. MERLoT uses diffusion maps to reduce the dimensionality of the expression vectors for each cell to a few components (typically between 2 and 20). Users can provide any low-dimensional manifold set of coordinates to MERLoT as input. **(B)** A scaffold tree is calculated given the low-dimensional manifold coordinates. **(C)** The scaffold tree is used to initialize a principal elastic tree, composed of k support nodes (default: 100), on which cells are assigned to the different branches of the tree. **(D)** Given a cell or tree node as the initial pseudotime to, pseudotime values propagate to the rest of cells/support nodes proportional to the distance along the tree that separates them from to. **(E)** Expression values from cells assigned to a given support node are averaged to provide it with an averaged expression profile for each gene. **(G)** A high-dimensional principal elastic tree is initialized with the connectivity from the low dimensional principal elastic tree plus the averaged expression values from the support nodes to impute smoothed gene gene expression data for each gene in the gene expression space. **(H)** MERLoT imputes the pseudotime-dependent expression profile of each gene along each branch in the tree.

To study gene expression changes along the different tree branches, MERLoT embeds the low-dimensional EPT structure into the high-dimensional gene expression space. Each tree support node in the low-dimensional space is mapped one-to-one to a tree support node in the gene expression space: We first assign each cell in the low-dimensional space to its nearest support node. Then, we initialize the corresponding support node in the gene expression space to the average gene expression level of all cells assigned to it (Fig. 1E), and we run the EPT algorithm again (Methods). In this way, we find the gene expression values of all the support nodes (Fig. 1F). These can be considered as pseudo-cells representing idealized cell differentiation paths.

The cells’ pseudotime values can also be refined in this step, since cells are reassigned to support nodes in the full multi-dimensional space. By combining the imputed expression values with the pseudotime assignments of the support nodes, MERLoT can reconstruct imputed pseudotime courses of gene expression profiles along the tree (Fig. 1G).

### Applying MERLoT to real datasets

We applied MERLoT on three real datasets with different degrees of lineage tree structure complexity (details in Methods): (1) scRNA-seq data (with Unique Molecular Identifiers, UMIs) for myeloid progenitor differentiation (2730 cells, 3460 genes) [16], embedded in DDRTree coordinates (Fig. 2A, D), (2) single-cell qPCR data for zygote to blastocyst differentiation (428 cells, 48 genes) [17], embedded in a diffusion map (Fig. 2B,E), and (3) scRNA-seq data (index-omics) for Haematopoietic Stem and Progenitor Cells (HSPCs) (1034 cells, 469 genes), using STEMNET coordinates [18]. The number of endpoints of the lineage trees, found by MERLoT in “auto” mode, are consistent with what has been reported in the original articles.

**Figure 2:**
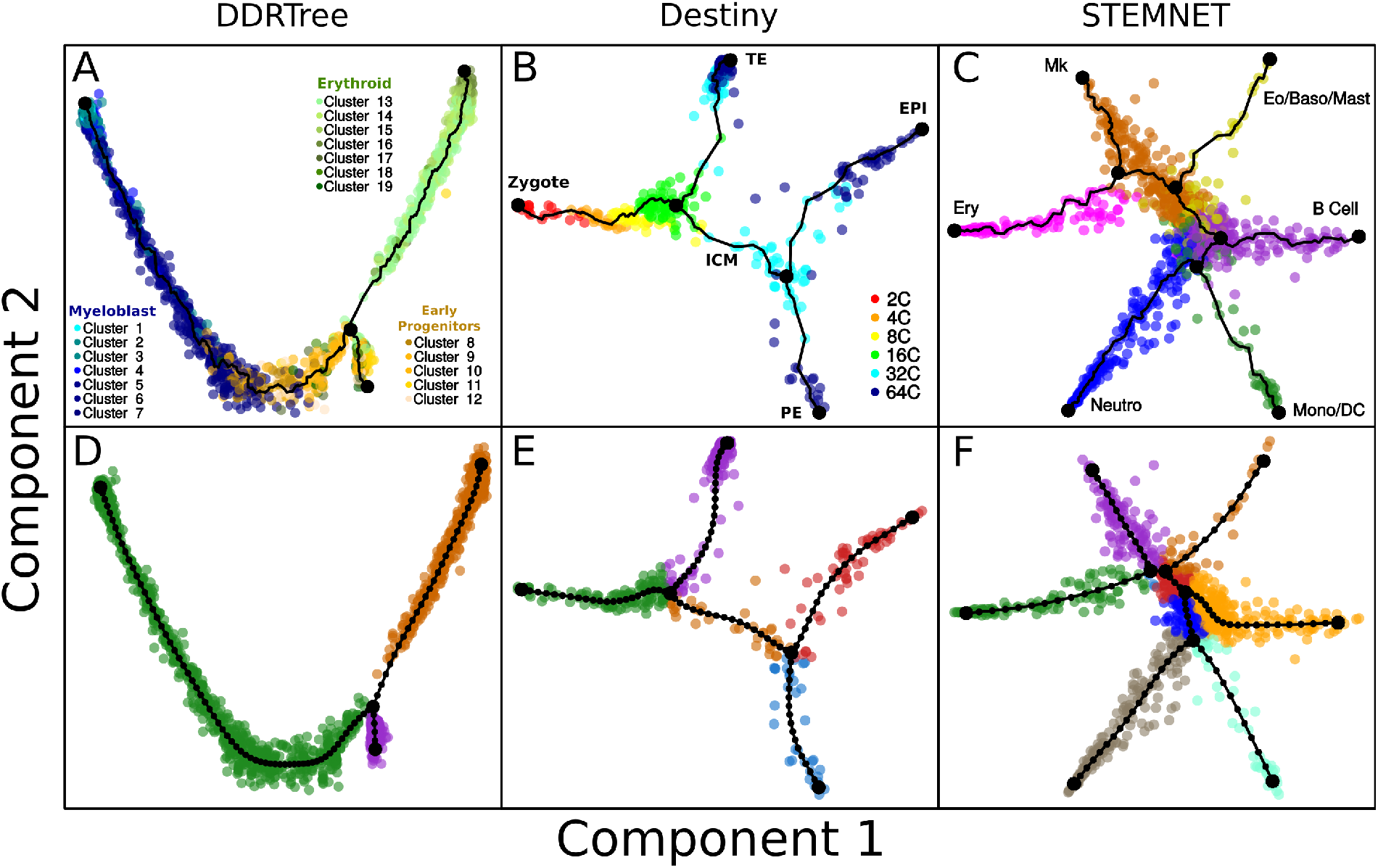
**MERLoT’s scaffold tree reconstructions:** in combination with **(A)** DDRTree coordinates for analyzing the data from Paul *et al*. [16]; **(B)** diffusion maps coordinates for analyzing the data from Guo *et al*. [17] (diffusion components 2 and 3 rotated around component 1 for better visualization of the data); **(C)** STEMNET coordinates for analyzing the data from Velten *et al*. [18]. Cells are colored according to cell type annotations provided by the authors of each dataset. **(D-F)** EPT reconstructions using the scaffold trees above as an initialization point. Cells are colored according to MERLoT’s branch assignments.

After the lineage tree reconstruction, each support node on the tree will be assigned a pseudotime value equal to the number of edges that separate it from the beginning of the differentiation, *t*_0_. This can be a tree endpoint (the Zygote branch in Fig. 2B) or an internal support node (for example a node in the red branch of Fig. 2E). Each cell will be assigned the pseudotime value of its closest tree node. In Fig. 3A, pseudotime values for the zygote to blastocyst dataset are shown. Because scRNA-seq data contain a lot of technical and biological noise [6], cells with similar pseudotime values may have large variations in gene expression. MERLoT imputes denoised gene expression profiles by embedding the reconstructed lineage tree into the original gene expression space.

**Figure 3:**
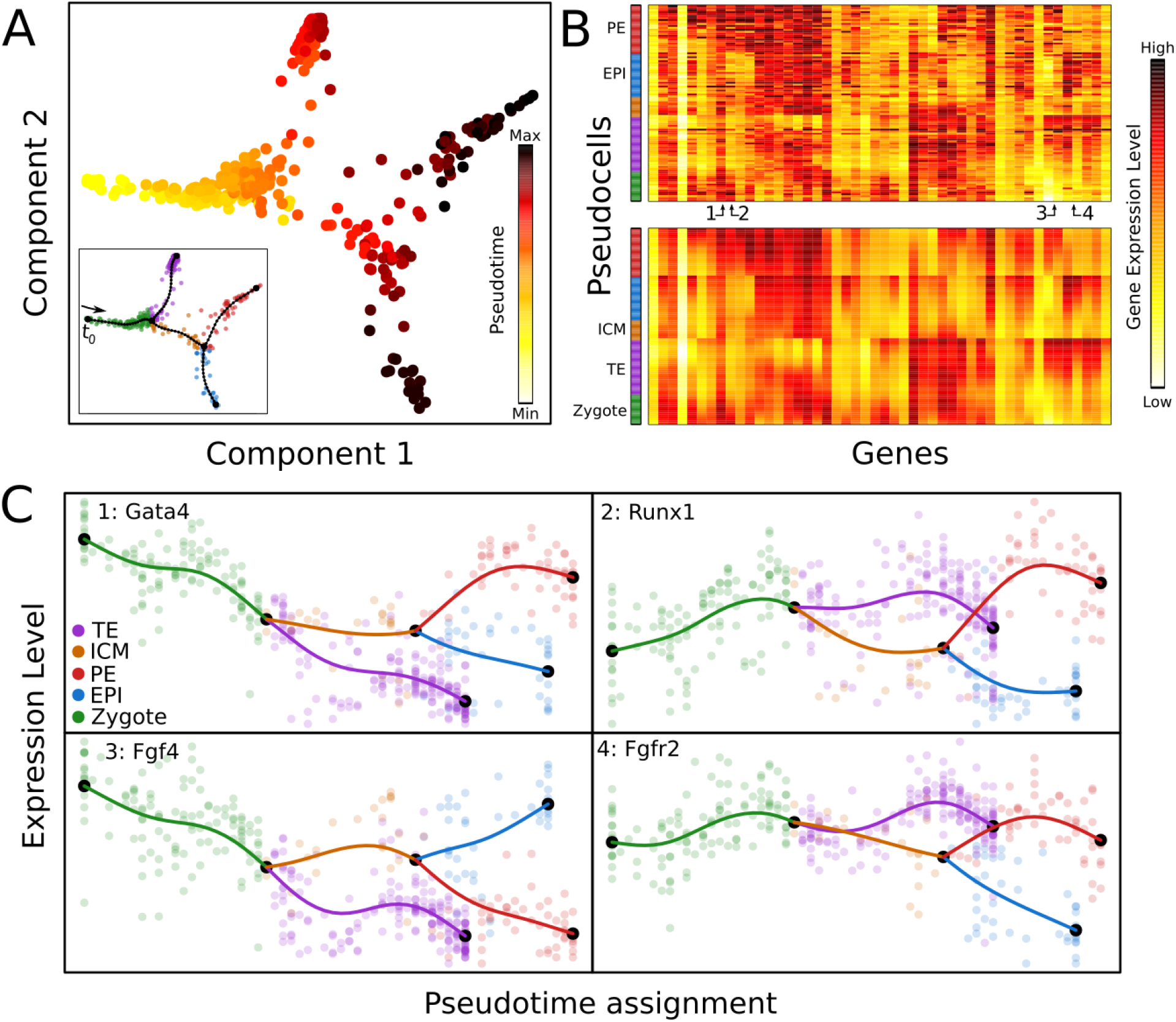
Pseudotime assignment and interpolation of gene expression profiles. **(A)** Pseudotime assignment in color code for cells from Figs 2B and 2E, taking the zygote state as *t*_0_. **(B)** Color-coded matrix of gene expression values for tree pseudo cells (rows) times genes (columns) before (top) and after (bottom) the gene expression space interpolation using the EPT. Pseudo cells are ordered according to their pseudotime and genes were hierarchically clustered. Numbers indicate specific genes shown in panel C. The color code in the bar on the left side of the heatmaps refers to pseudo cells branch assignments. **(C)** Gene expression profiles over pseudotime for 4 genes that are differentially expressed between the EPI and PE lineages. Semi-transparent circles represent the expression values of individual cells and solid lines correspond to MERLoT’s interpolations. Colors as in Fig. 2E.

This model-based interpolation results in denoised pseudotime courses of gene expression for the entire tree. As an example, Fig. 3C shows the expression profiles of four genes that are differentially expressed between the epiblast (EPI, in red) and the primitive endoderm (PE, in blue) cell lineages (Methods). Note how the expression values of the pseudo cells (solid lines) interpolate and smooth the noisy single-cell expression values (circles) even in regions with low cell density.

### Tree reconstruction performance assessment on synthetic data

In order to assess the quality of MERLoT’s lineage tree reconstruction we compared its performance to four tools with a similar approach to trajectory inference, namely unsupervised methods that produce branch assignments and assign a pseudotime to each cell: SLICER [19], Monocle2 [20], TSCAN [21], and Slingshot [22]. Since our focus was on correctly predicting cell labels and cell pseudotime, we needed data from complex topologies with known intrinsic developmental time.

For this purpose we developed PROSSTT (Methods, [11]), a software that can simulate scRNA-seq expression matrices with complex lineage tree structures. PROSSTT provides pseudotime and branch assignment labels for the cells, as well as branch connectivity information. Examples of diffusion maps for PROSSTT simulations and their lineage tree reconstructions with up to three bifurcations, performed by MERLoT, are shown in Fig. 4A. Additionally, we used Splatter (Methods, [12]), a suite to simulate cell populations with differential expression that can also be used to simulate branched topologies.

**Figure 4:**
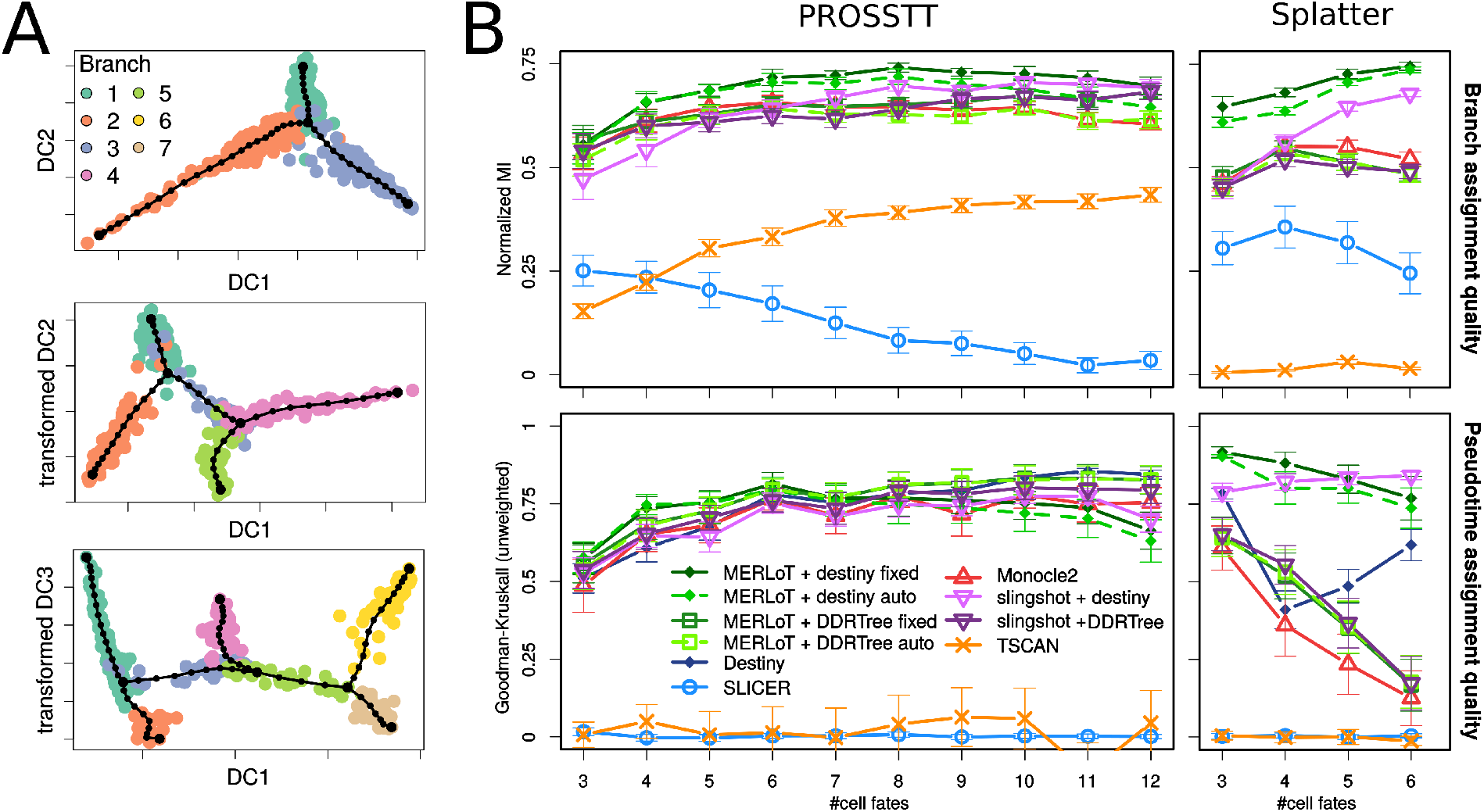
Simulated datasets and benchmarking. **(A)** Examples for diffusion map embeddings of PROSSTT simulations. From top to bottom: one, two, and three bifurcations. Cell colors the labeled branch assignments. The double bifurcation is plotted by rotating diffusion components 2 and 3 around component 1. The triple bifurcation is plotted by rotating diffusion components 3 and 4 around component 1. **(B)** Branch assignment comparison using Monocle2, SLICER, TSCAN, Slingshot and MERLoT using both PROSSTT (left) and Splatter (right) simulations. Slingshot and MERLoT are used in combination with DDRTree and diffusion map (Destiny) coordinates. **(C)** Pseudotime assignment comparison.

When dealing with complex topologies, methods usually perform better when they have access to more components in the dimensionality reduction step. However, visualizing hundreds or thousands of cells in multiple dimensions is challenging. MERLoT overcomes this limitation by using graph drawing techniques to visualize complex EPTs in two dimensions while still displaying cell annotation and density. In Fig. S3 we show reconstructed topologies for simulations with 6, 8, 10 and 12 different cell types.

We generated two simulated datasets. One contains 1000 simulations by PROSSTT and the other 400 simulations with Splatter. Each set contains subsets of 100 simulations. The PROSSTT set contains simulated trees with 3 to 12 different cell fates and 1 to 10 bifurcations respectively, while the Splatter set contains simulated trees with 3 to 6 cell fates and 1 to 4 bifurcations respectively. In the dimensionality reductions of datasets simulated by Splatter, branchpoints and endpoints often do not lie far enough from preceding tree segments. This makes it difficult for trajectory inference methods to correctly detect the tree structure, as they may connect tree segments that are in truth non-adjacent, thereby creating “short-circuits” (Methods, “Divergence Analysis” and Fig. S16). Because of this reason, we did not use higher order topologies simulated by Splatter for performance evaluation in the context of this benchmark. The results for the higher order bifurcations and more analysis on why they were left out are shown in Supplementary material (Figs. S4, S14, and S15.)

For all simulations, we predicted the lineage trees, assigned cells to branches, and calculated cell pseudotime values using the aforementioned tools. Since MERLoT and Slingshot work on a given set of manifold dimensions provided by the user, we used them in combination with diffusion maps (provided by the Destiny package) or DDRTree coordinates (provided by Monocle2). For simplicity we will refer to these combinations as MERLoT_Destiny, MERLoT_DDRTree, Slingshot_Destiny and Slingshot_DDRTree. Additionally, MERLoT can be used with and without providing the correct number of cell fates (“fixed” and “auto” modes). TSCAN and Slingshot do not provide a formal tree structure object (), so in order to evaluate their performance we had to implement wrappers for them (see Methods).

### Branch assignment quality

For branch assignment (Fig. 4B), we assessed the agreement between predicted and labeled predictions in the simulations, using the Normalized Mutual Information (NMI) (Fig. 4B) and other scoring measures: F1, Matthews Correlation Coefficient and the Jaccard Index (Methods and Fig. S4). Since NMI punishes splitting and merging clusters equally, it is not biased towards predictions having more or less branches than the simulations. For this reason, we consider it the best measure to assess cell assignment performance.

In the left side of Fig. 4B (top panel) we show the results for the PROSSTT set of simulations. SLICER and TSCAN perform worse than the rest of the tools. SLICER scores low mainly because its recursive branch assignment function crashes for many datasets or does not finish in less than 60 minutes, especially for complex topologies. TSCAN applies dimensionality reduction based on PCA, which is the main cause for its sub-optimal performance, given that the cells are all mixed up when projected in the PCA reduced space and hence cannot be correctly classified. MERLoT_Destiny performs best on almost all topologies except the simplest ones where they are very similar to other methods. MERLoT_DDRTree is slightly worse than Monocle2 for simple cases, however it improves as the topology complexity of the simulated trees increases, becoming better than Monocle2 for the most complex ones. In their article, Slingshot’s authors show trajectory inference examples using PCA coordinates which scored very poorly in our evaluation for the same cause TSCAN does. For this reason, we provided Slingshot with diffusion map and DDRTree coordinates as we did for MERLoT. Although for simple topologies Slingshot_DDRTree scores better than Slingshot_Destiny, the trend is reversed after inferring trees with more than 6 cell fates, with a similar performance compared to MERLoT_Destiny towards the end. From this, it seems that MERLoT works almost always better than the other methods, although Slingshot with diffusion maps ends up having a similar performance for the most complex topologies we simulated.

In order to better understand the trends about how the different tools perform, we analyzed the number of branches that each tool detects for the different sets of simulations (Fig. S8). TSCAN finds many more branches than the ones being simulated. Monocle2 and the MERLoT “auto” versions produce more branches than the simulated ones in the simplest topologies, but reverse the trend towards more complicated topologies at which they tend to underestimate the number of branches. Interestingly, in many cases, MERLoT using the “fixed” mode finds less branches than the simulated ones. This happens because when internal branches between bifurcations are too short, MERLoT can detect the same cell twice as a branchpoint. Hence, the tree topology is not completely binary but contains a trifurcation or higher order multifurcation, which decreases the total number of tree segments while maintaining the correct number of cell fates.

In the top right panel of Fig. 4B we can observe the results for the simulated set generated with Splatter. Since higher-order topologies presented short-circuits (Fig. S15), we only use differentiations with up to 4 bifurcations. Here we observe a wider spread for the performance of the different tools. MERLoT_Destiny is the best method for all topologies both using the “fixed” and “auto” modes. All tools that use the DDRTree embedding perform similarly well, with Monocle2 having a slight edge. SLICER and TSCAN still underperform, for the same reasons, while Slingshot_DDRTree also scores lower than Monocle2. Finally Slingshot_Destiny starts with a low performance, comparable to the one of Monocle2, but improves towards more complex topologies where it situates itself just behind MERLoT_Destiny and clearly above all other methods. For this set of simulations we observe in Fig. S8 that TSCAN produces extra branches compared to the simulated ones in a similar way it did for the PROSSTT simulations. This is also the case for Monocle2. For the rest of the tools there are no big differences, except for Slingshot_DDRTree, which detects less branches than in the PROSSTT simulations.

### Pseudotime assignment quality

In a multi-branched lineage tree, multiple trajectories exist between progenitors and differentiated cell fates. Pseudotime only imprints a partial ordering within each trajectory, and thus pseudotime values are not comparable between non-consecutive tree branches. We test pseudotime orderings on the cells that belong to the longest possible trajectory in terms of pseudotime steps in every simulation. We use the Goodman-Kruskal’s gamma (Fig. 4B, bottom) and other indices (see Methods and Fig. S6) as a measure of concordance between the true and predicted orderings along the longest trajectory.

As with the branch assignment assessment, there are no big differences in the performance of the various tools in the PROSSTT simulation set (left). MERLoT_Destiny is the best method initially, but falls off beyond 10 fates. Destiny becomes slightly better for more complex topologies, while Slingshot_Destiny steadily places in the middle of the pack. Slingshot_DDRTree mirrors the performance of Monocle, and MERLoT_DDRTree is the top tool for topologies with 5-8 bifurcations. Finally, the poor performance of TSCAN and SLICER at the branch assignment task in turn causes poor performance at pseudotime prediction.

In the Splatter set there is a general trend towards worse performance, something that can be explained by the growing number of short-circuits in the simulations. Monocle2 in particular seems to suffer considerably. MERLoT_Destiny outperforms the DDRTree methods, but also gets worse for more complex trajectories. Destiny itself improves over time after a dip in performance at the double bifurcation level. Slingshot_Destiny also shows some improvement for more complex topologies, however still remaining comparable to the performance of MERLoT_Destiny.

### Post lineage inference analysis

One of MERLoT’s assets is that it can reconstruct imputed gene expression profiles to study how a gene varies along the different paths in the lineage topology. These profiles interpolate and denoise the gene expression values of cells that are assigned to equivalent pseudotimes and facilitate the study/analysis of gene expression regulation, for example via detecting modules of genes that have correlated expression profiles. By using the imputed gene expression values recovered from MERLoT we can enhance downstream analysis like Gene Regulatory Network (GRN) reconstruction. We analysed the dataset in which Treutlein and coworkers studied the transdifferentiation process of Fibroblasts into Neurons [23]. Overexpression of the proneural pioneer factor *Ascl1* causes cells to exit the cell cycle and re-focus gene expression through distinct neural transcription factors. However, later on in the process a myogenic program competes with the neural one, producing undesired myocyte-like cells and lowering the efficiency of the direct reprogramming process.

We reconstructed the lineage tree from the data, which shows a single bifurcation (Fig. 5A). Then, we embedded the tree structure into the gene expression space and recovered the imputed gene expression values for the support nodes. We calculated the Pearson’s correlation coefficients between all pairs of genes using both the original gene expression values from the cells and the imputed ones from the tree support nodes. In Fig. 5B we show the Pearson’s correlation coefficient distributions for imputed and non-imputed gene expression values. While the non-imputed values concentrates most values between −0.5 and 0.5, the distribution of imputed values contains two subpopulations of genes close to −1 (highly anti-correlated) and 1 (highly correlated) separating them from the rest of weakly correlated genes. In Fig. 5C and 5D we show the gene expression values of S100a6, a gene differentially expressed in myocytes, and of Ap3b2, a gene differentially expressed in neurons, respectively.

**Figure 5:**
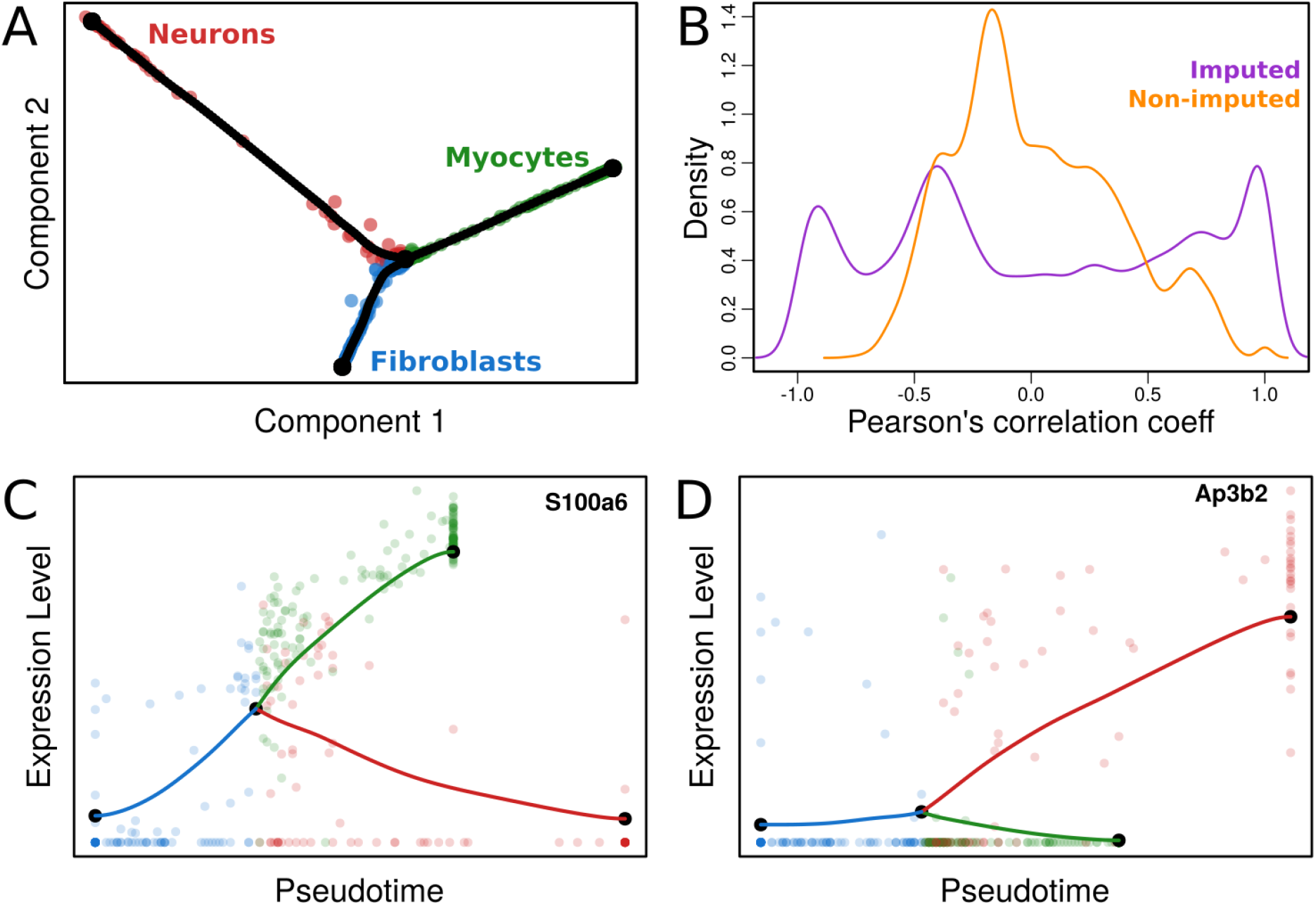
Lineage tree reconstruction of fibroblasts to neurons transdifferentiation. **(A)** Two-dimensional diffusion map embedding of cells together with the reconstructed lineage tree (tree support nodes shown in black). We observe a single bifurcated tree containing three branches corresponding to fibroblasts, neurons and myocytes. **(B)** Pairwise Pearson’s correlation coefficients for gene expression profiles using imputed (violet) and non-imputed (orange) values. **(C)** Pseudotime gene expression profile of differentially expressed gene in myocytes. **(D)** Pseudotime gene expression profile of a differentially expressed gene in neurons.

Once all pairwise gene expression correlations have been calculated, we built a graph where nodes correspond to genes and the pairwise Pearson’s correlation coefficients of imputed expression levels represent the weight of the edges connecting them. In Fig. 6 we observe the graph that results from applying a layout (see Methods) to distribute highly correlated genes (Pearson’s correlation coefficients greater than 0.9) in space. On this representation, close proximity corresponds to high correlation and vice versa.

**Figure 6:**
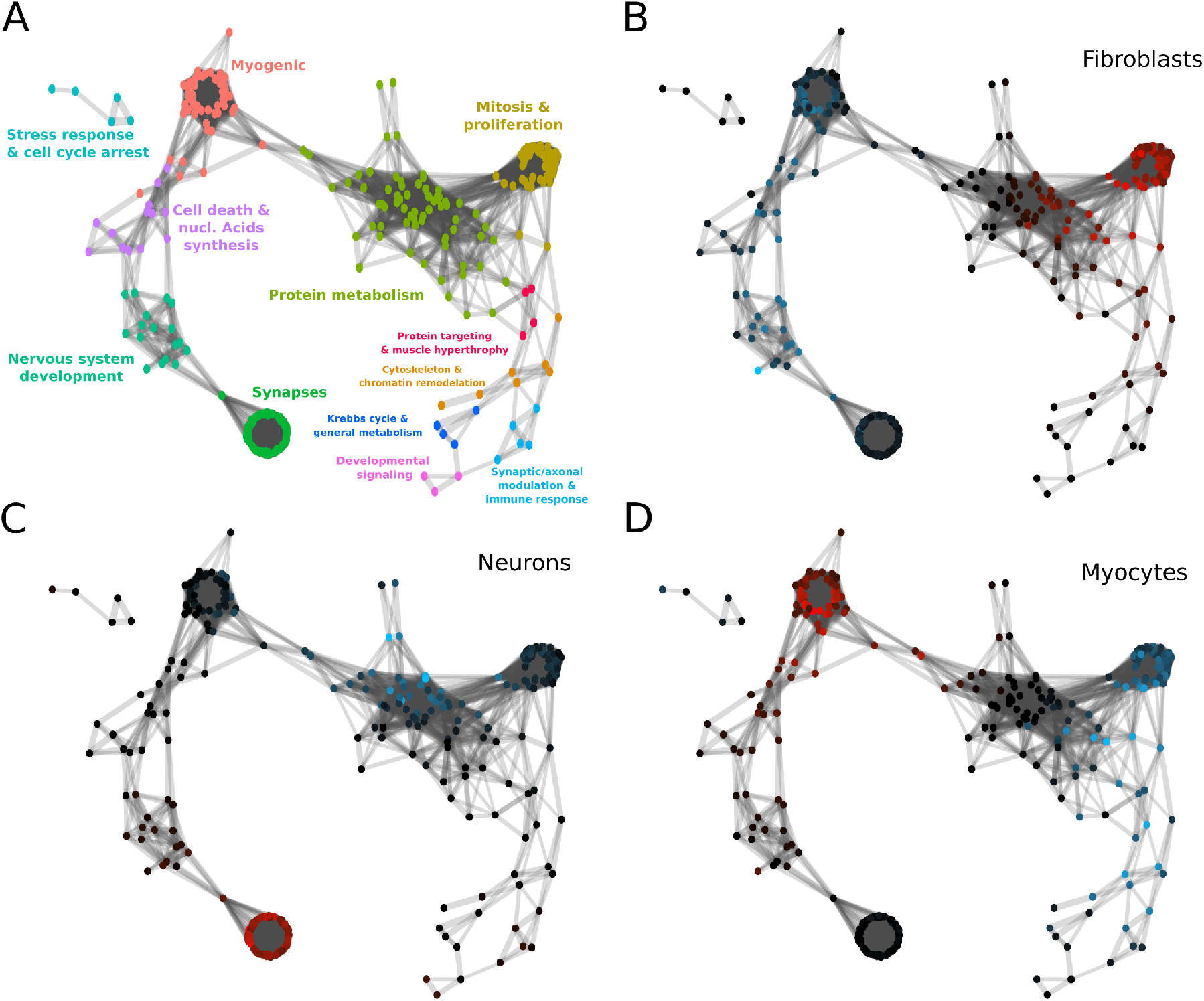
Reconstructed gene association network. **(A)** Gene clusters given the network connectivity were calculated and coloured. Enriched GO terms were retrieved for each cluster and a general label summarizing their main implications are assigned to each cluster. **(B)** Differentially expressed genes in the fibroblast branch. **(C)** Differentially expressed genes in the neurons branch. **(D)** Differentially expressed genes in the myocytes branch. Genes that are differentially expressed with an e-value *e* < 10^−3^ are colored according to the mean difference in expression, shades of blue indicating downregulation and shades of red upregulation. Intensity corresponds to log fold change of gene expression (see color scale).

For the quality of the reconstructed GRN, concentrating on highly-correlated genes and using imputed gene expression levels is crucial. A low correlation threshold will add noise to the GRN by including false positive interactions. Additionally, the noise in the non-imputed data obscures the similarities between the time-dependent expression of genes, resulting in low correlation values (Fig. 5B). This means that using a high correlation threshold will result in a GRN with few, small connected components, while a lower threshold will inevitably lead to densely connected “hairy ball” constructs that are difficult to interpret (Fig. S9-S11).

After obtaining the GRN, we clustered the network (see Methods) and recovered the gene ontology (GO) terms that were enriched in each cluster (see Methods). In Fig. 6A we show the reconstructed network coloured by cluster and labeled according to the keywords that best represent their enriched GO terms. We used MERLoT’s module for finding differentially expressed genes, both upregulated (red) and downregulated (blue), on each of the 3 subpopulations assigned to each branch that composes the lineage tree structure (see Methods). This was done for each branch, i.e, fibroblasts (Fig. 6B), neurons (Fig. 6C) and myocytes (Fig. 6D). Genes that are differentially upregulated in the fibroblasts branch mainly belong to the clusters enriched for GO terms related with “mitosis and proliferation” and “protein metabolism”. Downregulated genes in fibroblast cells mostly belong to clusters associated with the GO terms “myogenic”, “cell death and nucleic acids synthesis”, and “nervous system development”. We observe that for neurons, genes related to synaptic GO terms are upregulated while for myocytes the same happens for genes belonging to the myogenic cluster. Interestingly, downregulated genes on each branch belong to interconnected clusters. While neurons downregulate genes mostly associated with protein metabolism, myocytes downregulate genes associated with mitosis and proliferation, protein targeting and muscle hypertrophy, and cytoskeleton and chromatin remodeling.

## Conclusion

As the single-cell RNA sequencing field is becoming a mainstream technology, many datasets with highly complex underlying lineage trees will need to be analyzed. Here we have presented MERLoT, a tool to reconstruct complex lineage tree topologies in a more accurate way than other methods. We demonstrate this by applying MERLoT to various published datasets, but also by extensively testing its performance on a total of 1400 simulated datasets, produced by PROSSTT and Splatter. In this benchmark, MERLoT compares favorably to the state of the art in branch detection and pseudotime prediction using a variety of established performance indices.

Lineage tree reconstructions are not a final objective but rather a proxy to study changes in gene expression and understand the delicate regulation procedures that lead to organisms development, cellular differentiation, transdifferentiation and tissues regeneration. MERLoT simplifies and enhances downstream analysis in multiple ways. By utilizing an explicit tree structure, selecting subgroups of cells that belong to different tree segments or finding differentially expressed genes becomes straightforward. Apart from deriving the tree, MERLoT also imputes and interpolates gene expression, drastically reducing noise and alleviating the problem of gene dropout and hence enhancing downstream analysis. A recent publication described how diverse connective tissue cell types regenerated axolotl limbs after amputation by converging to the homogeneous transcriptional signatures of multipotent progenitor cells [24]. The authors used MERLoT to reconstruct the lineage tree, impute the gene expression values as a function of pseudotime, and study how gene expression levels changed among the different groups of cells in the process. Here, we have derived a gene regulatory network for a dataset representing the process of fibroblast to neuron transdifferentiation. We show that using MERLoT’s imputed expression values improves capture of gene-gene correlations, leading to more meaningful reconstructed gene regulatory networks.

While our benchmark demonstrated that MERLoT’s default approach leads to satisfactory results for a wide variety of topologies and expression matrices of various sizes, we are aware that when dealing with real data most methods don’t work out of the box. Currently, researchers rely on external, expert knowledge about the studied systems and manually optimize strategies on every step of the process (gene selection filtering, cell quality filtering, different manifold embeddings, and tuning of parameters related to all of these). MERLoT is flexible enough to allow supervision at different parts of the analysis pipeline, while providing default strategies that are robust enough to be used for exploratory analysis.

Single-cell RNA sequencing allows for the study of time-dependent processes in unprecedented detail. With the help of tools like MERLoT, we can overcome the high noise and non-linearities in the data, reconstruct the cellular lineage trees, and follow the change of gene expression over developmental time. Such time course gene expression profiles pave the way for the reconstruction of gene regulatory networks and eventually their quantitative modelling, which will profoundly advance our understanding of developmental processes.

## Methods

### 1 Lineage Tree Reconstruction by MERLoT

Given an expression matrix with *N* cells as rows and *G* genes as columns, a manifold embedding can be performed using techniques like Diffusion Maps or DDRTree embedding. Subsequently, informative dimensions can be kept in order to reduce dimensionality. Once cells are embedded into the low dimensional space, MERLoT can perform a lineage tree reconstruction following three steps which will be detailed in the following sections: (1) Calculating a Scaffold Tree with the location of endpoints, branchpoints and their connectivity. (2) Smoothing of the Scaffold Tree by using an Elastic Principal Tree (EPT) in the low dimensional manifold. (3) Embedding the EPT into the high-dimensional gene expression space and assigning pseudotime values to the cells.

#### Scaffold Tree Reconstruction

##### Matrix of shortest paths between cells

We used the Dijkstra’s algorithm as implemented in the `csgraph` module from the `scipy` python library to calculate the shortest paths between all pairs of cells *i* and *j* based on their squared Euclidean distance 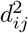. We use 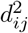 instead of *d_ij_* given that in a fully connected graph the shortest path between two nodes, based on *d_ij_*, corresponds to the direct path between them. The `csgraph shortest_path` function returns the length *D_ij_* of the shortest path that connects cells *i* and *j*. We extended the function to also return the number of cells *S_ij_* on the shortest path connecting cells *i* and *j*. The modified code for `csgraph` is available at github.com/soedinglab/csgraph_mod.

##### Endpoints search

MERLoT does not require users to define a starting point in order to reconstruct the tree topology. The first two endpoints in the tree correspond to the pair of cells *k, l* that maximize the number of cells *S_kl_* on the shortest path between them (Fig. S1A): (*k, l*) = argmax{*S_kl_* : 1 ≤ *k* ≤ *l* ≤ *N*}. In case of ties, the pair of cells with the longest shortest path distance will be selected.

The next endpoint to be selected will correspond to the cell *n* that maximizes 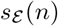, the number of cells being added to the scaffold tree structure. To compute 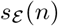 we note that the new endpoint *n* must branch off from an internal node (cell) of the tree whose next-neighbour nodes must lie on the path of two already selected endpoints *k* and *l*. Therefore, the increase in number of cells is 0.5 times the minimum of the cells between *k* and *l via n* minus the cells between *k* and *l*, minimized over all pairs of endpoints (*k, l*) in the set of already selected endpoints 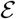.

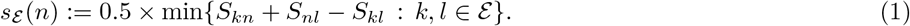

##### Stop criterion for endpoint search

In auto mode every time a new endpoint is selected we evaluate if 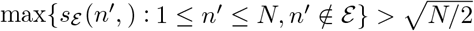 holds true. Otherwise, we calculate the branchpoints and tree connectivity for the endpoints in 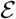, including *n*, using the methodology explained in the next subsection. After this, all cells are mapped to their closest branch. If the branch added by the selected *n* endpoint contains more than 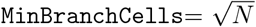 cells mapped to it, the branch is kept in the tree scaffold structure and the endpoints search is repeated. Otherwise, the endpoint search terminates and *n* is discarded as endpoint. The `MinBranchCells` threshold can be modified by the user. Alternatively, instead of using a stop criterium, users can set the number of endpoints that are aimed to be found (fixed mode) regardless of the branch lengths.

#### Branchpoints search and tree connectivity definition

Once endpoints are found, we apply the in combination with a heuristic criterion to find the cells that best represent the branchpoints in the tree structure. By doing this, we also detect the tree connectivity among endpoints and branchpoints Given a set of endpoint and branchpoint cells we iterate the following steps: (1) Use the Neighbour Joining (NJ) criterion [14] to select the pair of (*k, l*) cells that will be joined *via* a branchpoint next. 2) Find the cell *m* that best represents the branchpoint between *k* and *l* and add the *l* − *m* and *m* − *k* edges to the tree (Fig. S2).

Let 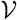 be the set of yet unprocessed endpoint and branchpoint nodes of the tree. We initialize 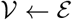 with the endpoint set 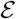 determined in the previous subsection.

1. Given the matrix *D_kl_* of shortest path lengths between nodes in 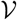 (section 1.1), the NJ criterion allows us to pick two nodes *k, l* in 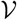 that are guaranteed to be next neighbours and therefore can be linked via a single branchpoint. The nodes to be joined are chosen such that 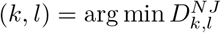 where 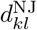 is the neighbour-joining distance,

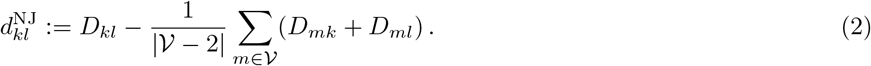
2. We determine the branchpoint cell *m* as the one that minimizes the sum of distances to nodes *k, l* and the mean distances to all other nodes included in 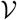,

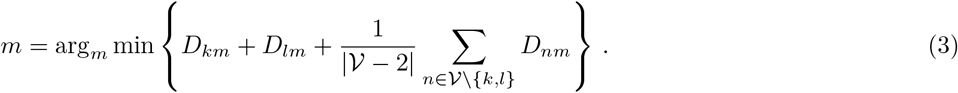 Next, *k* and *l* are removed from 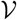 and the new branchpoint *m* is added to 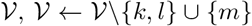. Also, the edges *l* − *m* and *m* − *k* are added to the tree (Fig. S2).

We iterate (1) and (2) until 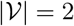, which means no further branchpoints exist. After termination, we have determined the tree topology with its 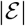 endpoints and 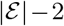 branchpoints, each represented by a cell. The same cell can be detected more than once as a branchpoint and hence bifurcations with higher orders than binary ones are possible.

#### Elastic Principal Tree in the Low Dimensional Manifold

To produce smoother, more homogeneously interpolated lineage trees MERLoT uses the Elastic Principal Trees (EPT) algorithm [15] as implemented in the ElPiGraph.R module (https://github.com/Albluca/ElPiGraph.R and [25]). The EPT algorithm is used to approximate the distribution of cells in a given space with a tree structure composed of *k* nodes. Direct application of the EPT algorithm is unstable as it often returns trees that are manifestly far from the global optimum, e.g. wrong number of endpoints or grossly misplaced branchpoints). We initialize the EPT with the coordinates of a set of *l* nodes consisting of the endpoints and branchpoints in the scaffold tree and their connectivity. Further nodes are then added by the EPT algorithm by iterative bisection of edges until it reaches the specified number of *k* support nodes. This procedure cannot modify the number of endpoints and branchpoints specified at the initialization step.

In every iteration an energy potential defined as follows is minimized:

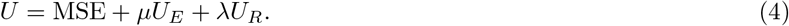

MSE is the sum of squared distances of cells to their closest tree support nodes. *U_E_* and *U_R_* are two regularization terms that ensure that we learn trees with regularly spaced points and with edges without kinks, respectively.

We performed a grid search around the EPT default values in order to optimize *μ* and λ and visually examined the reconstructed EPTs on the datasets shown in Fig. 2. Using *k* = 100, we obtained *μ*_0_ = 0.0025 and λ_0_ = 0.8 · 10^−9^. If the number of nodes used for calculating the EPT is changed, *μ* and λ are adjusted according to *μ* = (*k* − 1)*μ*_0_ and λ = (*k* − 2)^3^λ_0_. All reconstructions in our benchmark were performed with the standard function using *k* = 100. However, for some particular topologies *μ* and λ might need to be tuned in order to produce optimal results, in particular if *k* is increased a lot. Alternatively, MERLoT can bisect the edges in a given EPT, by additional nodes producing a new EPT with almost 2*k* support nodes. Note that these are special cases.

#### Elastic Principal Tree Embedding into the Gene Expression Space

First, cells are assigned to their closest support node. Next, every node is assigned to the average expression profile of all cells assigned to it. If a given node is not the closest node to any cell in the set, a null vector is assigned to it. In this way, we translate the positions of cells in the low-dimensional manifold space to approximate positions in the full gene expression space by constructing *k* pseudocells with averaged values for each gene coordinate.

Finally, the EPT algorithm is initialized with the average expression profiles and a list of edges representing their connectivity in the low dimensional EPT to calculate an EPT in the high-dimensional gene expression space.

##### Pseudotime Assignment

Pseudotime is a quantitative measure of the progress of a cell through a biological process [26]. Given the reconstruction of a lineage tree by MERLoT, cells can be assigned pseudotime values as a function of the number of edges along the structure that separate them from the initial point of the process. MERLoT automatically sets the initial pseudotime, to, to one of the first two detected endpoints. Users can also set *t*_0_ to any endpoint, branchpoint or to any individual cell. In the latter case, the closest node to that cell will be assigned as *t*_0_ and the pseudotime values for the other nodes will be assigned as before. After a pseudotime value is assigned to each support node, cells will take the pseudotime value from their closest node in the tree. MERLoT can calculate pseudotime in both the low-dimensional manifold space and in the high-dimensional gene expression space.

### 2 Benchmark on Synthetic Datasets

#### 2.1 Simulating count data of branching processes with PROSSTT

To evaluate method performance on reconstructing lineage tree structures we used our tool PROSSTT [11] to simulate 10 × 100 scRNA-seq datasets with topologies with between 1 and 10 branchpoints (3-12 endpoints).

PROSSTT generates a simulated scRNA-seq dataset in four steps: (1) it generates a tree (number and length of branches, connectivity), (2) it simulates average gene expression levels *μ_g_*(*t,b*) (pseudotime *t*, branch *b*), (3) it samples points in the tree (*t, b*) (4) it retrieves *μ_g_*(*t,b*) for each sampled point and draws UMI counts from a negative binomial distribution

##### 1. Generate tree

We sample the number of genes from a discrete uniform distribution between 100 and 1000. These are typical numbers left in real datasets after filtering out uninformative genes.

Each tree segment in every simulation had a pseudotime length of 50, corresponding to 50 cells on the branch (in homogeneous sampling model). This length allows the expression programs to diffuse enough to be distinct from each other. Starting at 2 bifurcations, alternative segment connectivity possibilities become available (Fig. S7)). We chose the lineage tree topology randomly for every simulation.

##### 2. Simulate average gene expression along tree

PROSSTT models relative gene expression as a linear mixture of a small number of expression programs. For each tree segment, we simulate the time evolution of expression programs as a random walk with momentum term. Each simulation uses *K* = 5*b* + *u* expression programs, where *b* is the number of branchpoints and *u* is drawn from a uniform integer distribution 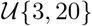. Each program contributes to the expression of every gene, with weights drawn from a gamma distribution (shape parameter *a* scales inversely with number of programs, rate parameter *b* is 1). A scaling factor (library size) was sampled for each cell and multiplied to the average gene expression values. We used a log-normal distribution with *μ* = 0 and *σ* = 0.7.

##### 3. Sample cells from tree

PROSSTT can sample cells in the tree according to a given density function. Here we used a uniform density and drew 50 × *b* cells, where *b* is the number of branches.

##### 4. Simulate UMI counts

We simulate unique molecular identifier (UMI) counts using a negative binomial distribution. Following [27] and [28], we make the variance 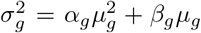 depend on the expected expression *μ_g_* as 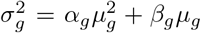. The *α_g_* and *β_g_* values were sampled from log-normal distributions with *μ_a_g__* = 0.2, 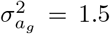 and *μ_b_g__* = 2, 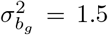 respectively. These values are typical for single-cell RNA sequencing UMI counts. For more information about default parameters choices and the algorithm, please refer to [11].

We provide the scripts used to create the simulations (https://github.com/soedinglab/merlot-scripts) as well as the simulations themselves (http://wwwuser.gwdg.de/~compbiol/merlot/).

#### 2.2 Simulating count data of branching processes with Splatter

In order to validate the results produced by the PROSSTT benchmark, we used Splatter [12] to simulate another 10 × 100 scRNA-seq datasets with topologies with between 1 and 10 branchpoints (3-12 endpoints). Splatter takes a slightly different approach to the simulation of lineage trees (“paths” in the terminology of Splatter). It is a software primarily designed to simulate populations of cells with differential expression between them. In order to simulate a lineage tree, it simulates populations with differential expression at each successive waypoint of the lineage tree (between start and first branchpoint, between successive branchpoints, between branchpoints and endpoints), and then simulates how gene expression changes from one waypoint to the other.

For parameter selection, we kept default parameters as far as the count model and the generation of average gene expression values were concerned (parameters controlling mean, library size, expression outlier, biological coefficient of variation). The rest of the simulation parameters were picked to mirror those in the PROSSTT simulations:

- **Global parameters:** the number of genes was picked from the same discrete uniform distribution 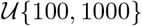. A random seed was set and included to ensure reproducibility.
- **Batch parameters:** The number of batch cells (substitutes total number of cells if only one batch is present, as is the case here) was set to 100 × b, where b is the number of branches. This is twice the number sampled for PROSSTT simulations, which increased the robustness of the dimensionality reduction. The other batch parameters were left to their default values.
- **Group parameters:** The number of groups was set to the number of branches, and the occurrence probability of each group was set to 1/*b*.
- **Differential expression parameters:** Since we were only interested in simulating informative genes, we set the probability of differential expression per gene to 1. All other parameters were left at their default values.
- **Differentiation path parameters:** The topology of the lineage tree was input as a vector of originating points per branch (`path.from` parameter). Branch lengths were set to 50 via `path.length`.

#### 2.3 Assessing Methods Performance

Tree reconstruction tools need to succeed at two intertwined tasks: to arrange all cells according to their internal developmental time, while also detecting and separating different branches of the tree. Ideally, one would evaluate algorithm performance on both tasks at once. However, we were not able to find such a measure and so evaluate branch assignment and pseudotime prediction separately.

##### Branch assignment

We treated branch assignment evaluation as a clustering problem. We can consider all cells mapped to a given branch of a tree as a cluster and compare the set of labels produced by PROSSTT with those that are predicted by each algorithm.

We used four measures to assess method performance: the F1 measure, the Matthews Correlation Coefficient, the Jaccard Index and Adjusted Mutual Information (called Normalized Mutual Information, or NMI, here) (Fig. S4), which all produce similar results. In our opinion the NMI is best for assessment of branch assignment, since it captures the amount of information present in the original clustering that was recovered by the prediction (values between 0 and 1). It corrects the effect of agreement solely due to chance between clusterings by using a hypergeometric background distribution and punishes overbranching and merging branches almost equally [29]. Given the predicted and real cluster assignments *U* and *V* with *R* and *C* clusters respectively, NMI is defined as

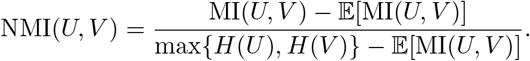

where *H*(*U*) is the entropy of *U*, MI(*U, V*) is the mutual information of *U, V* and 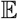 denotes the expectation value under the null model of independent *U* and *V*.

##### Pseudotime prediction

Evaluating the performance of the different methods consists of quantifying the degree of agreement between the true/labeled and the predicted orderings provided by the different algorithms. Pseudotime only establishes a partial ordering on cells and no absolute time. Therefore, cells on branches not passed through one after the other cannot be compared. We find the longest path in the tree (from the root to a leaf) and compare the predicted pseudotime with the simulated one for the cells on this path.

In this sense, pseudotime prediction assessment is a comparison of ordered sequences, and we follow the suggestions of [30] by using the Goodman-Kruskall index and the Kendall index (weighted and unweighted). All four indices count how many pairs of cells have been ordered correctly (*S*_+_) or incorrectly (*S*_−_) and produce similar results for the benchmark. The unweighted Goodman-Kruskall index is the simplest approach: (*S*_+_ − *S*_−_)/(*S*_+_ + *S*_−_).

#### 2.4 Benchmarked Methods and Parameters

##### Monocle2

Monocle2 (version 2.6.1) uses a reverse graph embedding (RGE) technique called DDRTree [31] to create a lineage tree and map points from the low-dimensional embedding of the tree to the original gene space. Based on the vignette for (`monocle_2.4.0`), we used the `negbinomial()` expression family for the data and the proposed defaults for lower detection limit (1), minimum expression (`detectGenes` function, 0.1), mean expression threshold (gene ordering, 0.5), and empirical dispersion threshold (2 · *dispersion_fit*).

Generally, a successful dimensionality reduction will capture a topology with *B* bifurcations in its first *d* = *B* + 1 dimensions. While Monocle2 runs DDRTree by default on two dimensions, in our benchmarks we used *B* + 1 dimensions, as it performed better. DDRTree did not always return *d* dimensions; if fewer dimensions were provided we used the maximum possible number.

Monocle2 assigns a branch identity to each cell in the `State` column of the phenotypic data table (`pData`). It calculates pseudotime as distance from one of the endpoints of the lineage tree it produces. In order to pick the correct endpoint, we checked if the cell with simulated pseudotime *t*_0_ = 0 was in an outer branch; if true, the corresponding endpoint was assigned *t*_0_. In the rare event that it was placed in an inner branch, we used Monocle’s chosen starting `State` for pseudotime calculation.

##### SLICER

SLICER uses locally linear embedding to perform dimensionality reduction and uses a neighbour graph to order cells according to their distance from a user-specified starting cell (pseudotime prediction). It uses geodesic entropy to recursively detect branches.

We used SLICER version 0.2.0 (commit cb1be8a) by following the instructions that accompany the software on its github page (https://github.com/jw156605/SLICER/). We used the software’s gene selection process as-is and used the selected genes to determine the best *k* value for the LLE, with a *k*_min_ value of 5. Much like with Monocle2, using *d* = *B* + 1 LLE dimensions (over the default 2) for a dataset with *B* bifurcations improved performance. We used the same *k* value for the creation of the low-dimensional k-nearest neighbour graph as we did for LLE. For every simulation, we used the cell with labeled pseudotime *t*_0_ = 0 as the starting point of the `cell_order` function, which predicts pseudotime for each cell. Finally, we used the same start point for the branch assignment step (`assign_branches`). This step very often failed to execute; these cases were assigned the worst possible score of each branch assignment measure. The issue was reported to the authors (https://github.com/jw156605/SLICER/issues/7) but until the time of this writing was not resolved.

##### Destiny

Destiny produces a dimensionality reduction and a pseudotime prediction based on it. We used the destiny diffusion map space as input for MERLoT, and benchmarked destiny’s Diffusion Pseudotime (DPT). We used destiny (version 2.6.1) as described in the Bioconductor vignette. First we normalized the input data by correcting for library size and then log-transformed them. The only free parameter is the number *k* of nearest neighbours. By default, destiny uses a heuristic to determine it. For a dataset with *B* bifurcations we gave destiny *d* = *B* + 1 dimensions of the diffusion map to determine pseudotime, following the same reasoning as with Monocle2 and SLICER. We retrieved pseudotime predictions from destiny by calling the `DPT` function on the diffusion map object and then using the dimension which corresponded to diffusion pseudotime distances from the cell with pseudotime *t*_0_ = 0.

##### MERLoT

We ran different flavors of MERLoT on embeddings from different dimensionality reduction tools:

1. **MERLoT + diffusion maps:** destiny was run to produce diffusion maps for MERLoT. The same protocol as in section 2.4 was used, except for the selection of free parameter *k*, which was done by a simple optimization. For a simulated lineage tree with *B* bifurcations we tested values of *k* between 5 and 100 and kept the *k* value that maximized the drop-off after the *d* + 1’th eigenvalue of the diffusion map, where *d* = *B* + 1.
2. **MERLoT + DDRTree:** Monocle2 was run in order to produce DDRTRee coordinates. The number of coordinates to be used to reconstruct the lineage tree was selected as described in section 2.4.

Each of these two options was combined with two different ways in which MERLoT finds the number of endpoints in the dataset.

1. **MERLoT auto:** MERLoT is run without the specification of the number of endpoints in the tree. MERLoT will use its internal branch length heuristic to determine new branches. The algorithm stops finding new endpoints when the new branch aimed to be included in the tree structure does not contain more than 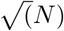 number of cells, with N being the total number of cells in the dataset.
2. **MERLoT fixed:** MERLoT is run with a specified number of endpoints (tree leaves) to be found. MERLoT will ignore its internal branch length heuristic and will keep searching for branches until it reaches the specified number of endpoints regardless of how many cells are mapped to them. All 2 × 2 combinations were tested (MERLoT on diffusion maps auto/fixed and MERLoT on DDRTree coordinates auto/fixed).

#### 2.5 Divergence analysis

While benchmarking method performance on data simulated with Splatter, we noticed that multiple methods did not perform according to expectations. The issue was particularly obvious in the evaluation of pseudotime prediction (Fig. S13D), where Monocle2 sank to the level of random predictions, and MERLoT, which, even though it excelled at branch assignment (Fig. S13B), dropped off to quite low accuracy for large numbers of bifurcations. Additionally, TSCAN, which in the PROSSTT benchmark proved to be competent in branch assignment (Fig. S13A), returned completely nonsensical predictions for data simulated by Splatter.

As these methods have all been applied successfully on real data, we chose to examine the simulations. By visual inspection of the manifold embeddings we observed that the manifold embeddings of Splatter simulations often presented “short-circuits”, where parts of the lineage tree seemed to fold back to preceding tree segments, effectively creating cycles. While branches were often separated correctly (i.e. the cell clustering was correct), they were connected in wrong ways, decreasing the pseudotime prediction accuracy.

Since the manifold embeddings are a projection that aims to retain the most important dynamics in the data, we hypothesized that the reason for these short-circuits was that the offending branches did not diverge enough, or even converged towards previous tree segments. In terms of gene expression, this means that differentiating phenotypes (captured transcriptomes) were either not different enough or that they converged towards preceding phenotypes.

To test this hypothesis, we measured the distance of each waypoint (endpoint or branchpoint) from the origin, and normalized it by the average branch length in the path that led to it. The calculations described below were performed on the simulated gene expression data, after normalization for library size and log-transformation (see scripts `divergence_euclidean.R and divergence_diffmaps.R`).

As explained in Fig. S14, we retrieved the cells with minimum and maximum pseudotime in each branch (start cells and end cells respectively), calculated their pairwise distances, and defined their average *d_b_* as the length of branch *b*. Next, we took the cells with globally minimum pseudotime and calculated their distances from all end cells. This is the minimum direct distance 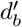 of each branch *b* from the origin. We normalized each 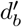 with 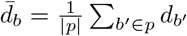, where *p* is the path [*b*_0_, *b*_1_, …*b*] from the origin branch *b*_0_ to branch *b*. Effectively, this yields the distance from the origin measured in average branch lengths. Pooling the normalized distances from all paths of equal length (especially since all branches have the same length in the PROSSTT and Splatter simulations) shows how much the change in expression values correlates with pseudotime.

The divergence curve of PROSSTT (Fig. S15A) shows what we expected: monotonic growth (i.e. longer paths are on average further away from the origin) and values above 1, indicating that paths with two branches or more consistently end outside a one-branch-length radius from the origin, something that reduces the possibility of wrong assignments from the methods.

On the contrary, the divergence curve of Splatter (Fig. S15B) stays almost completely below 1 and even shows a slightly negative slope. This means that expression profiles of cells in later differentiation stages dont move further from the origin with increasing pseudotime length. We believe this happens because Splatter does not include co-regulation in its differentiation simulation model. This leads to the differences between branches being completely random, and while this may work to separate two diverging branches from each other, it does not seem to yield realistic diverging cell lineage trees.

As a control, we performed the divergence analysis in the diffusion maps created by destiny for the benchmark (panels G,H in Fig. S15). Diffusion distance was proposed as a measure of cell similarity in the original paper [32], and was the most effective of the embeddings used in this study. We see the same trends as in gene expression space; the divergence curve of PROSSTT has positive slope and is consistently above 1, while the Splatter curve has a (clear) negative slope and stays below 1.

After quantifying divergence in both PROSSTT and Splatter simulations, we conclude that the simulations produced by Splatter have inherent characteristics that prevent algorithms that try to find a global structure (like MERLoT and monocle) from reaching their full potential. Consequently, we decided to only use simpler topologies for the Splatter benchmark (up to 4 bifurcations), where the impact of short-circuits was less dominant.

### 3 Datasets

#### Myeloid progenitors differentiation (Paul, et al. Cell, 2015 [16])

This dataset was produced applying massively parallel single-cell RNA-seq (MARS-seq) which uses unique molecular identifiers (UMIs). After filtering those cells that passed the quality checks and selecting a set of informative genes we obtained a matrix of 2730 cells and 3459 genes. Originally the authors reported 3461 informative genes with some of them being incorrectly formatted as dates, e.g 5-Mar, 4-Sep. We were able to correct the IDs of most of them to valid GeneIDs except two (IDs: 7-Sep and 2-Mar) which were excluded from the analysis. The script for calculating the DDRTree coordinates using Monocle2 and the subsequent lineage tree calculation (Fig. 2A and Fig. 2D) is available at: https://github.com/soedinglab/merlot/tree/master/inst/example/ExamplePaul2015.R.

#### Mouse zygote to blastocyst (Guo et. al, Developmental Cell, 2010 [17])

The dataset was produced by the Biomark RT-qPCR system and contains Ct values for 48 genes measured in 442 mouse embryonic stem cells at 7 different developmental time points, from the zygote to blastocyst [17]. The data was cleaned and normalized by following the vignette from the Destiny package. A total number of 428 cells and 48 genes were kept in the final expression matrix. A diffusion map was calculated using Destiny and the first three diffusion coordinates were used to calculate the lineage tree. We rotated the cells and tree nodes coordinates around the first axis in order to produce a two-dimensional representation of the data and improve visualization. The script for calculating the diffusion maps and the subsequent lineage tree calculation (Fig. 2B and Fig. 2E) is available at: https://github.com/soedinglab/merlot/tree/master/inst/example/ExampleGuo2010.R.

#### Haematopoietic Stem and Progenitor Cells (HSPCs) (Velten et. al, Nature Cell Biology, 2017 [18])

The scRNA-seq data was generated samples taken from two donor individuals. The authors used both smart-seq2.HSC (individual 1) and QUARTZ-seq (individual 2), and all findings were systematically compared between individuals. For obtaining the processed data we followed the vignette for the STEMNET software available as part of its R package. From that analysis we obtained the normalized data, the cell types labels and the STEMNET coordinates. The script for calculating the the STEMNET coordinates and the subsequent lineage tree calculation (Fig. 2C and Fig. 2F) is available at: https://github.com/soedinglab/merlot/tree/master/inst/example/ExampleVelten2017.R.

#### Differentially expressed genes detection

After a linear tree reconstruction has been performed, MERLoT can easily find groups of genes being differentially expressed among different groups of cells. If two groups of cells are provided, e.g cells assigned to two branches in the tree (Fig. 3C)), MERLoT performs a Kruskall-Wallis rank sum test to evaluate which genes in the full expression matrix are differentially expressed on them. If a single subpopulation of cells is provided, the comparison is made against the rest of cells in the data. The entire list of genes is given as output, ordered by the test p-values results. Also, e-values are provided by multiplying the p-values by the number of *G* genes being tested.

#### Regulatory Network Reconstruction

We performed a Gene Regulatory Network (GRN) reconstruction for the fibroblasts to neurons transdifferentiation dataset from the Treutlein group [23]. We reconstructed the lineage tree, reconstructed the GRN, performed gene clustering and differential gene expression analysis. The script for performing this analysis is available at: https://github.com/soedinglab/merlot/tree/master/inst/example/

##### Network construction

Given the expression profile of a gene in all cells (non-imputed values) or in the tree support nodes (imputed values) we create a matrix that contains the pairwise Pearson’s correlation coefficient between all pairs of genes. Given this matrix we can use the R package *igraph* (http://igraph.org/r/) by defining a threshold to decide which Pearson’s correlation coefficients to be include as weights for the edges in the graph.

##### Cluster Analysis

We clustered the genes on the GRN by applying the *walktrap* algorithm as implemented in the *igraph* package. This algorithm partitions a graph into modules based on the logic than short random walks tend to stay within a module. Any module with less than a specified lower boundary of nodes is dissolved and those genes are considered as unclustered. We set the minimum number of genes to 3.

##### GO term enrichment analysis

To help determine the function of network defined clusters, a module for gene ontology (GO) term enrichment was implemented. For this purpose the R package topGO was used. GO term enrichment was computed by the Fisher exact test with the gene set of the data set as the gene universe. Gene ontology tables per cluster were retrieved and representative keyboards were selected by cluster.

##### Differentially expressed genes

We detected the differentially expressed genes at every branch of the reconstructed lineage tree using the `branch_differential_ex` function from MERLoT. In shades of red, upregulated genes. In shades of blue, downregulated genes. All genes are ranked according to the e-value obtained from the “Kruskall-Wallis rank sum test” applied to test whether they are differentially expressed or not. Genes with e-val < 10^−3^ are considered to be differentially expressed. Genes are colored according to the mean difference in expression between the two compared sets of cells (i.e selected branch and rest of the tree for each case). Blue colors indicate downregulation and red colors upregulation. Intensity corresponds to log fold change of gene expression. Genes that are not significantly differentially expressed are colored in black.

## Data availability

The 10 simulation sets with 100 simulated differentiations each are available at http://wwwuser.gwdg.de/~compbiol/merlot/. The code necessary to run the benchmark on the simulations as well as instructions about how to set up a similar benchmark are available at https://github.com/soedinglab/merlot-scripts. Formatted expression data for the three datasets in Fig. 2 are available at: https://github.com/soedinglab/merlot/tree/master/inst/example/

